# Pathway maps enable straightforward yet customized and semi-automated yet insightful analyses of omics data

**DOI:** 10.1101/404525

**Authors:** Steffen Möller, Israel Barrantes, Robert Jaster, Larissa Henze, Hugo Murua Escobar, Christian Junghanss, Oliver Hakenberg, Marc-André Weber, Bernd J. Krause, Olaf Wolkenhauer, Saleh Ibrahim, Rüdiger Köhling, Falko Lange, Uwe Walter, Mohamed Hamed, Axel Kowald, Georg Fuellen

## Abstract

To explore the molecular processes underlying some biological theme of interest based on public data, gene lists are used herein as input for the construction of annotated pathway maps, employing Cytoscape apps, and then high-throughput (“omics”) gene expression data are overlaid onto these maps. Seeded with a published set of marker genes of the *senescence-associated secretory phenotype* and the genes of the *cellular senescence* KEGG pathway, a gene/protein interaction network and annotated clusters (a “pathway map”) of cellular senescence are derived. The map can be amended, by adding some application-specific genes, and overlaid with gene expression data describing cellular senescence of fibroblasts and with disease-related gene expression data associated with prostate and pancreatic cancer, and with ischemic stroke, allowing insights into the role of cellular senescence in disease. Some gene expression data are derived from the “Biomarker Benchmark repository”. The pathway map approach can be followed in principle for any biological theme of interest, fostering much-needed independence from the investigator-biased expert networks usually used for overlaying gene expression data.

## Introduction

High-throughput data do not usually yield biological or medical insight just by themselves. Enrichment analyses are arguably the most popular way of generating insight, but they do not usually consider the mechanistic details of gene-protein interaction and regulation, as pathways and interaction networks do. Thus, there is a need for a flexible, easy to follow route to detailed insight, which does not limit the researcher to a specific fixed set of genes to begin with (like, KEGG pathways), nor to the expert knowledge as it is ingrained in a pathway database. Moreover, such a route to detailed insight should be simple and straightforward. Also, the route to detailed insight should be as robust as possible, and this property is not easy to fulfill. In this work, robustness specifically refers to a high degree of stability of the clusters into which the network unfolds, with respect to modifications of the input gene list. Without such clustering, we are left with interaction networks in the form of unstructured “hairballs” that enable fewer insights^1^. However, if we shy away from the commonplace utilization of more or less immutable expert pathways (from KEGG, WikiPathways, Ingenuity, etc.), we need to obtain the pathway or network interaction information based on other sources, which are necessarily non-expert-curated interaction data. These are available in large amounts, but these are also inherently noisy and based on a mix of experimental or computational source interactions generated in a variety of contexts. Thus, we must expect automated clustering to lack robustness. Nevertheless, by an appropriate choice of interaction data, we here demonstrate that it is possible to generate pathway maps based on gene lists in an automated fashion. Moreover, we show that these pathway maps can be sufficiently stable such that small perturbations of the input gene list, e.g. the addition of a few genes, do not trigger sweeping changes in the pathway map, even if we insist on *non-overlapping* clusters, for easy comprehension and visualization. Using the MCL clustering and annotations based on wordcloud-assisted processing of GO gene annotations as provided by Cytoscape apps (for details see below), in this paper we assemble a plausible pathway map that is describing cellular senescence in a highly unbiased fashion. We furthermore amend the pathway map by adding a few genes based on application-specific interest. For senescence- and disease-related data, we then show how the pathway maps make it easier to extract insights from high-throughput gene expression datasets.

The focus of our interest for establishing such an automated workflow is on cellular senescence. 50 years ago it was discovered that human diploid fibroblasts have a finite replicative potential in culture after which the cells enter a state of irreversible replicative arrest^2^. Today it is clear that in addition also various types of stress, like reactive oxygen species or DNA damaging agents, can induce cellular senescence, suggesting that it is a special stress response state of the cell^3^. Transcriptional changes include an up-regulation of tumor suppressor and anti-apoptotic genes and a down-regulation of cell-cycle promoting genes. In addition, senescent cells secrete an inflammatory mix of cytokines, growth factors and matrix metalloproteinases, which form the *senescence-associated secretory phenotype* (SASP). This paracrine signaling has a range of negative effects involving tissue remodeling, aging and tumorigenesis. Molecular identification of senescent cells is not trivial, since the senescent state induced by different triggers in different tissues is heterogeneous^4^. Still, key markers are a large and flat cell morphology, a senescence-associated form of β-galactosidase, and expression of tumor suppressors such as CDKN1A (p16^INK4A^). Senescent cells are involved in the initiation and progression of various diseases. Although cellular senescence generally acts as a tumor suppressor mechanism, it can also promote cancerogenesis and fibrosis via the SASP^5^. Such (antagonistic) pleiotropy includes fibrotic processes important for wound healing^6^ and in the liver^7^. Cellular senescence is also implicated in diseases such as cancer, stroke, atherosclerosis, osteoarthritis and metabolic disorders^5^. A causal relationship is supported by studies that showed that transplanting senescent cells into young animals caused physical dysfunction^8^ and removing senescent cells increases health and lifespan^8-10^.

Prostate and pancreatic cancer are on opposite ends in terms of survival prospects at time of diagnosis. Prostate cancer is a heterogenous disease ranging from well differentiated and hardly progressive low-grade cancer to highly aggressive and life-threatening high grade disease. In the United States, prostate cancer that is local or regional at the time of diagnosis has a 5-year survival rate of nearly 100%, while those with distant metastases have a 5-year survival rate of 29%^11^. In contrast, in the less than 20% of cases of pancreatic adenocarcinoma with a diagnosis of localized small cancerous growth (less than 2 cm in Stage T1), about 20% of Americans survive to five years^12^. Cellular senescence can suppress both prostate and pancreatic cancer, and cancerous proliferation in general, but it also triggers tumor progression by the SASP^13,14^. Also, cellular senescence contributes to atherosclerosis and thromboembolism, and after the ischemic stroke it can attenuate recovery^15,16,17^. Moreover, cancer and stroke are linked by components of the SASP, specifically PAI1 (aka SERPINE1)^18^. Patients with pancreatic cancer show an especially high incidence of thromboembolic complication^16^. In the following, we specifically explore such molecular commonalities by mapping high-throughput data to senescence-related pathway maps.

## Results

### Construction and exploration of senescence pathway maps

In the following, we will first explore a canonical map (based on SASP-related and cellular senescence genes), overlaying public high-throughput data that were specifically generated to characterize cellular senescence. We will then explore the canonical map with disease-related data. We also add genes of interest to explore specific senescence pathway maps for specific disease applications.

Applying GeneMANIA^19^ and AutoAnnotate^20^ to the 189 SASP and cellular senescence genes^4,21^ (see Suppl. Table 1 in Suppl. File 1), with default parameters (except for the limitation of interaction data to co-localization, genetic and protein interactions, see *Methods*), we obtained the pathway map of Fig. 1 and Table 1. GeneMANIA added 20 closely interacting genes towards a total of 209 genes. In Fig. 1, the nodes in the pathway are colored using three gene expression data sets of Ras-induced senescence^22,23,4^, as described below. The clusters from Fig. 1 that include more than two genes are presented in Table 1, where the numbering matches the one in the figure, starting top-left. All clusters and full lists of genes are provided in Suppl. Table 2 in Suppl. File 1. The Cytoscape file is provided in Suppl. File 2. Finally, the pathway map can be explored interactively at http://functional.domains/senescence/.

**Fig. 1:**
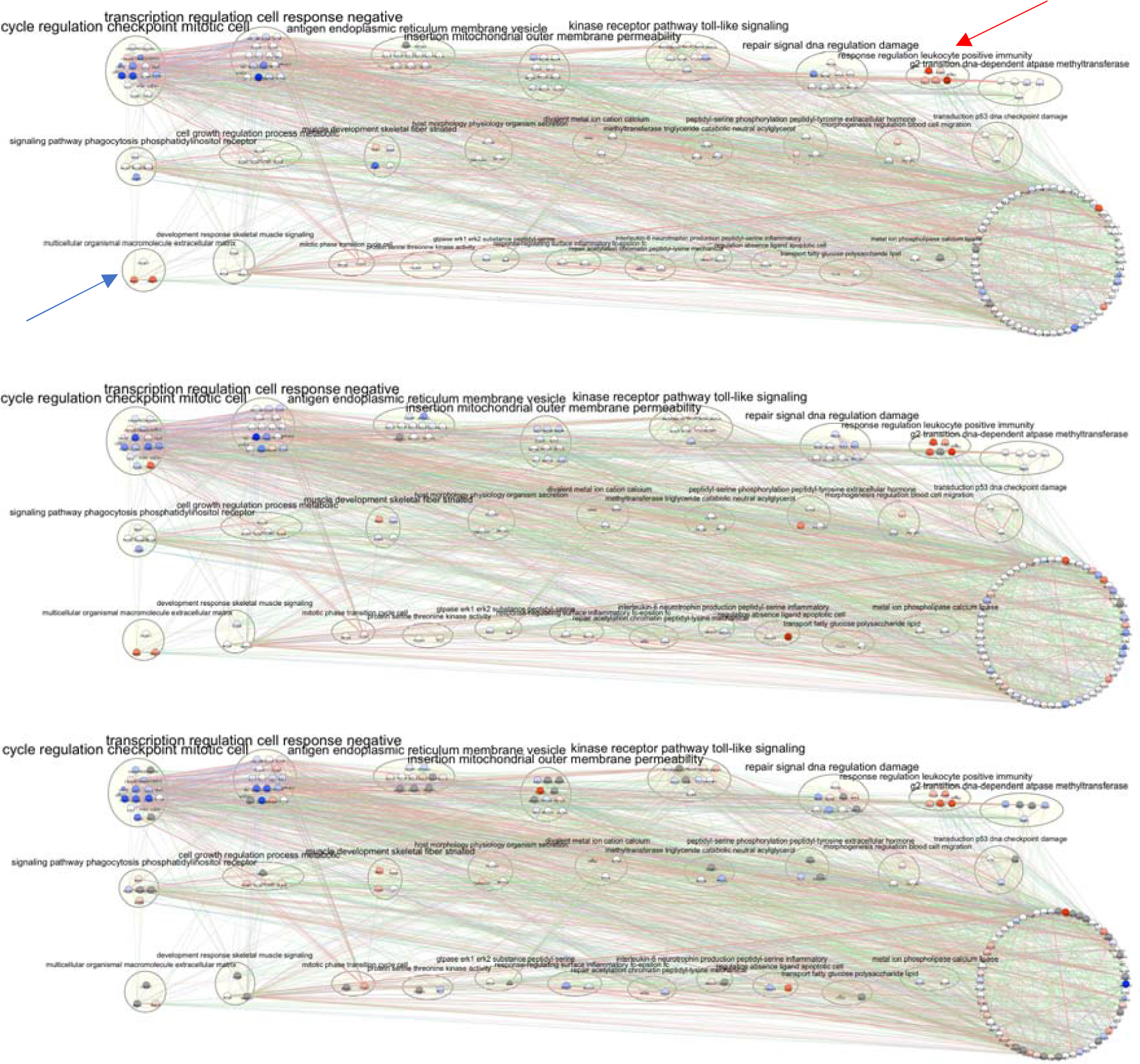
Canonical senescence pathway map, based on the SASP and the KEGG cellular senescence genes. The same pathway map is shown with annotations from three different experiments. The size of a gene node is proportional to its GeneMANIA score, which indicates the relevance of the gene with respect to the original list of genes to which GeneMANIA, based on the network data, added another 20 genes. Genes upregulated in senescence (GSE19899, ref^22^, top, GSE61130, ref^23^, middle, E-MTAB-5403, ref^4^, bottom) are shown in red, downregulated genes are shown in blue, and grey denotes genes for which no expression values were available. Clusters with genes known for their association with the SASP are indicated by an arrow. The color of an edge refers to the source of the edge in the underlying network, that is physical interactions (red), co-localization (blue), and genetic interactions (green). The thickness of an edge is proportional to its GeneMANIA “normalized max weight”, based on the network data. The pathway maps can be explored interactively at http://functional.domains/senescence.

**Table 1:**
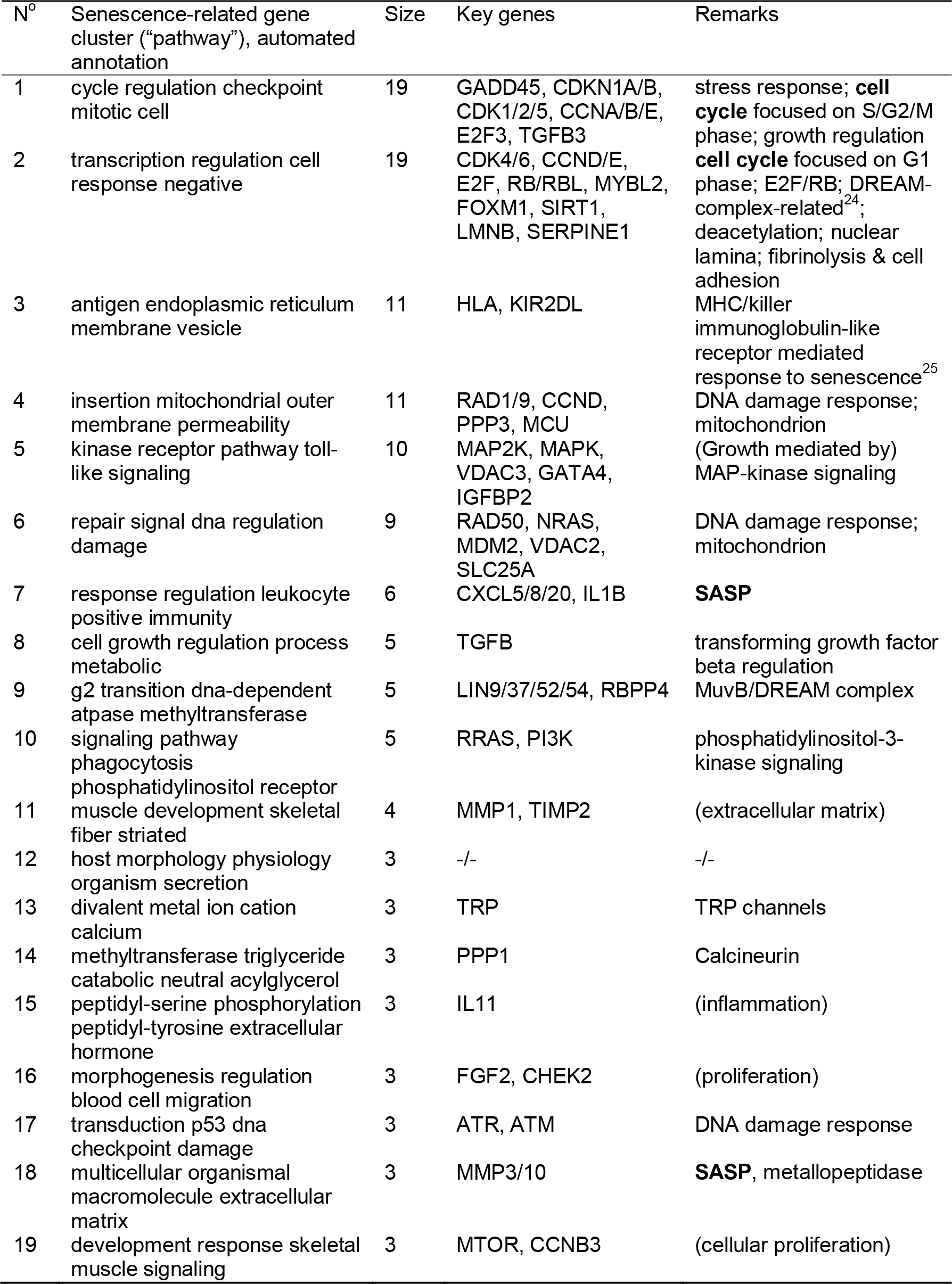
Senescence-related clusters/pathways of Fig. 1, sorted by size. Suppl. Table 2 in Suppl. File 1 provides a full list of clusters (including clusters of size 2), and the full list of genes per cluster.

In a bird’s-eye view, the canonical map of Fig. 1 consists of 28 clusters. The two largest clusters/pathways on the top left feature most of the cell-cycle genes, including CDKNs (cyclin-dependent kinase inhibitors), CDKs (cyclin-dependent kinases) and CCNs (cyclins). The SASP is mostly featured in the 6-gene cluster near the top right (cluster 7, red arrow), and in the 3-gene cluster bottom-left (cluster 18, blue arrow). The bottom-right circular structure includes all genes not assigned to any cluster. Table 1 provides a list of the larger clusters; we will refer to this detailed breakdown of clusters/pathways in the presentations that follow. By construction, the clustering rests on nothing else but co-localization, protein and genetic interaction data, and the annotation of these clusters rests on nothing else but the GO annotations of the genes in each cluster. Thus, the pathway map required no specific expert intervention except for the individual layout of the clusters, which placed the genes roughly in an upstream/downstream fashion (see *Discussion)*.

### Exploring microarray data on cellular senescence

The gene expression data on Ras-induced senescence of IMR90 fibroblasts displayed in Fig. 1 (top) were published as part of a study of the role of the Rb (retinoblastoma) protein family in senescence^22^. The authors reported that a “disruption of a p21-mediated cell-cycle checkpoint” could be due to loss of the tumor suppressor Rb that usually attenuates cellular proliferation. Accordingly, in the situation of senescence as described in Fig. 1 (top), we find that p21 (CDKN1A) is the only strongly upregulated gene in the cluster/pathway 1, and the most strongly downregulated genes are the cyclins (CCNs) A2 and B1, reflecting that the S/G2/M phases are most affected (Vermeulen et al, 2003). In cluster/pathway 2, we find that other parts of the cell cycle (G1 phase) are not affected as much. Most prominently, downregulation of Lamin B is observed here. Lamin B happens to be allocated to cluster/pathway 2, even though its downregulation is a general feature of cellular senescence. The two clusters/pathways most closely associated to the SASP (pathways 7 and 18, next-to-top-right and bottom-left of the map, red and blue arrow) feature the expected upregulation of their members, in particular of IL1B and of cytokines, and of matrix metalloproteinases 3 and 10. In cluster/pathway 11, MMP1 is found upregulated as well, and its inhibitor TIMP1 is downregulated. Ras-induced senescence is reflected in the map by upregulation of HRAS, which is however part of the circular structure of genes not allocated to any cluster/pathway. Finally, the members of the Rb family^22^ only feature a negligible fold-change, suggesting that their regulation is not mediated by the amount of transcript, but likely by phosphorylation, as is also the case for the CDKs. The E2F transcription factors do not feature much fold-change either; their downregulation in case of senescence is plausible though, as they are considered tumor drivers that are inhibited by the Rb proteins. Finally, CCNE1 (cyclin E1) as the key target of Rb is upregulated and this unexpected observation is discussed extensively in Chicas, et al. ^22^.

### Exploring RNA-Seq data on cellular senescence

For comparison, we overlaid the RNA-Seq based expression data of Herranz, et al. ^23^, which were used to study Ras-induced senescence in IMR90 fibroblasts, see Fig. 1 (middle). For this dataset, the change in gene expression (log fold change, logFC) could be calculated for 205 of the 209 genes in the senescence pathway map; this number is the largest one available of all public datasets considered by Hernandez-Segura, et al. ^4^. We note the high concordance between Fig. 1 top and middle; in cluster/pathway 1, the Herranz dataset is different only in that CXCL2 is upregulated, while in cluster 2, SERPINE1 is up-regulated instead of down-regulated, but Lamin B is downregulated as in the first IMR90 senescence dataset. The SASP in clusters 7 and 18 is also upregulated, and the SASP-related antagonism of MMP1 and TIMP1 in cluster 11 is visible. The SASP factor CSF2^26^ (cluster 25) is upregulated much stronger here; in fact it is the strongest-upregulated gene in Fig. 1 (middle) but it is not further discussed in Herranz, et al. ^23^. Also, IL11 (cluster 15) is strongly upregulated.

We also explored the datasets generated by Hernandez-Segura, et al. ^4^, along which our list of SASP genes was published. We focused on the HCA2 human foreskin fibroblast data of radiation-induced senescence (disregarding keratinocyte and melanocyte data) to allow for the most meaningful comparison with the other fibroblast data we investigated, using the 20-day post radiation data set that reflects cell-cycle arrest best according to the paper. This data set also has the highest number of genes (of the canonical senescence pathway map) for which fold changes are available (160 out of 209). Reassuringly, the gene expression landscape (Fig. 1, bottom) closely resembles the one observed before, e.g. for clusters/pathways 1 and 2 (except that CDK6 and CCNE1 feature higher expression). In cluster 4, CCND2 is upregulated strongly. The SASP is again upregulated, and in cluster 25, CSF2 is upregulated here as it is in the Herranz dataset (Fig. 1, compare middle to bottom).

### Exploration of cancer, stroke and mitochondrial dysfunction data

#### Prostate cancer

We utilized the “Biomarker Benchmark repository” of Golightly, et al. ^27^, based, for prostate cancer, in turn on Erho, et al. ^28^ (GSE46691), and we used the repository’s gene expression data in R^29^ to derive log fold change data describing prostate cancer disease progression (logFC from Gleason grade 7 to 8-10, which is prognostic for metastasis, as suggested by the repository’s “Prognostic__Metastasis_Analysis.txt” file). For mapping to the senescence pathway map, we inverted the colors in this case, as disease progression towards metastasis is opposite to tumor-suppression by cellular senescence. Reassuringly, SASP genes are then turning red (that is, up in senescence, down in metastasis) in clusters 7 and 18 (and also in cluster 1, affecting CXCL2), and cell cycle genes in clusters 1 and 2 are turning blue (Fig. 2, top). The secretion of CXCL2 as a pro-inflammatory cytokine was found when investigating the stromal-epithelial interactions in the early stages of prostate cancer, in an in-vitro setting^30^.

**Fig. 2:**
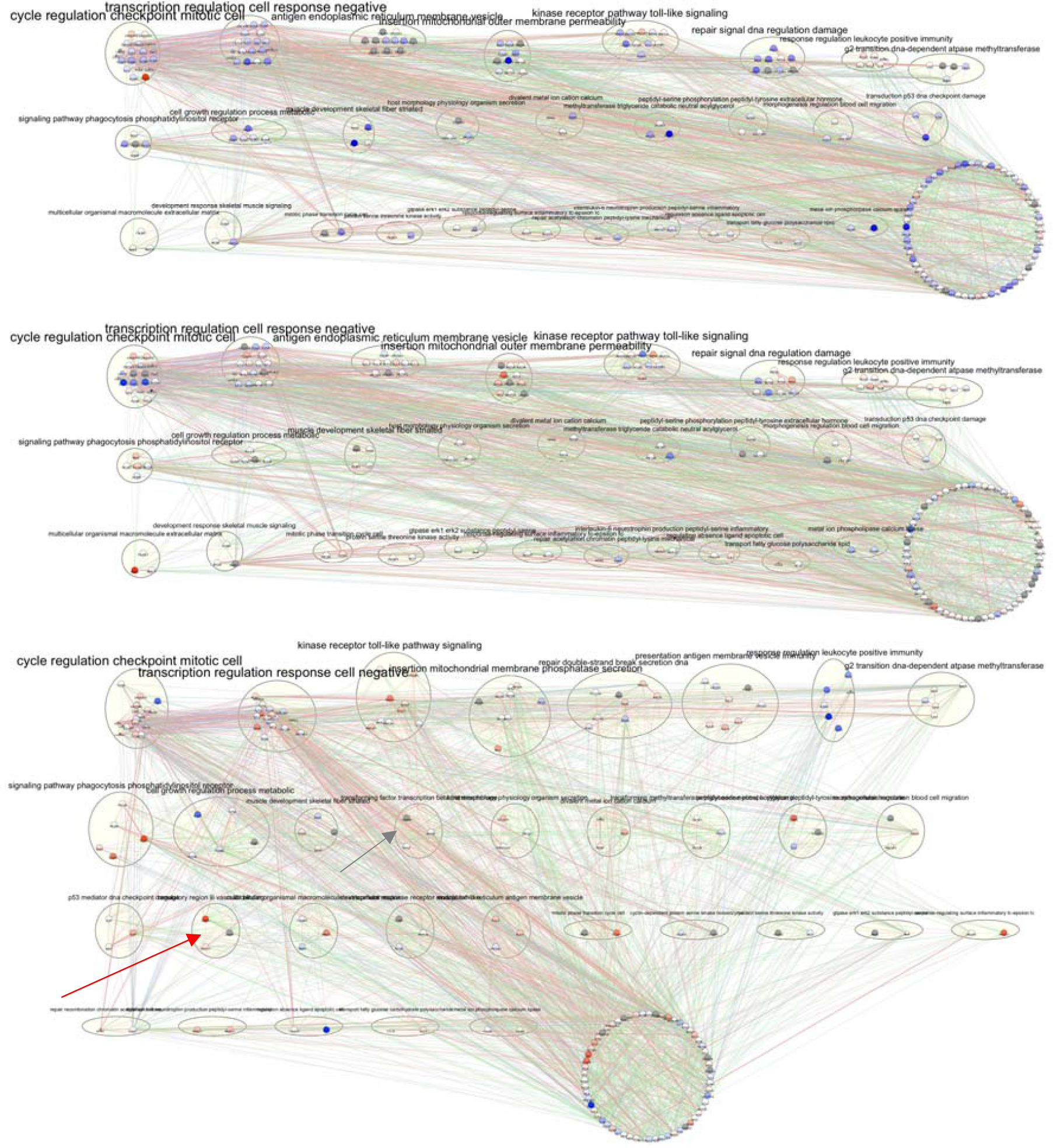
Senescence in disease, prostate and pancreatic cancer. The canonical senescence pathway map (top and middle), based on the SASP and the KEGG cellular senescence genes, and an amended one (bottom), adding two genes of interest and omitting any manual layout. Genes upregulated in disease (GSE46691 Erho, et al. ^28^, top, GSE81368, ref^31^, middle, GSE28155, ref^32^, bottom) are shown in red, downregulated genes are shown in blue, and grey denotes genes for which no expression values are available. See Fig. 1 for further explanations.

#### Pancreatic cancer

Overlaying gene expression data of carcinoma-associated fibroblasts (GSE81368)^31^ onto the canonical senescence pathway map, in the first pathway (cluster 1) we observe downregulation of most cell-cycle genes, specifically of cyclins A2, B1 and B2, matching the observation of a reduced number of S-phase cells reported by the authors (see Fig. 2, middle). The upstream stress signaling by GADD45 is upregulated, as is the downstream SASP-related factor CXCL2. In cluster 2, which is more closely connected to the G1 phase, there is no consistent pattern. SASP genes in clusters 7 and 18 are upregulated, most prominently CXCL8 (also known as IL8) and MMP3, both as noted in the paper. Still, cyclin D variants, in form of CCND1 (in cluster 20), CCND2 (in cluster 4) and CCND3 (in cluster 2), are upregulated, suggesting some G1-phase activity. Next, we overlaid gene expression data describing that senescence driven by KDM6B, a tumor-suppressing mediator of KRAS-induced senescence, attenuated aggressiveness of PDAC cells (GSE28155)^32^, onto an amended senescence pathway map, following our map construction recipe except for adding KDM6B and its downstream target CEBPA (also known as C/EBPα) to the list of input genes. The amended senescence pathway map in Fig. 2 bottom was thus generated *de novo*, and it is displayed without any manual layout. The clustering, however, is essentially the same as in the canonical pathway map. Again, clusters 1 and 2 feature most of the cell cycle genes. While gene expression of KDM6B (grey arrow) was not measured, CEBPA (red arrow) is downregulated as expected. However, the cell cycle gene featured in the paper, CDKN2A/p16, is not differentially expressed as would be expected, the other cell cycle genes are regulated in an inconsistent fashion, and the SASP genes are unexpectedly downregulated (cluster 7).

#### Stroke

Overlaying the differential data provided by comparing ischemic stroke patients to controls^33^ (GSE22255) reveals that senescence-associated processes can indeed be found in peripheral blood mononuclear cells (PBMCs) after stroke. Specifically, in cluster 1, CDKN1A is upregulated, and, correspondingly, CCNA2 is downregulated, though most cell cycle genes are regulated in no consistent fashion. The SASP genes in cluster 7 are upregulated, as is interleukin 6 (IL6; found in the circular structure bottom right). IL6 is not only an important part of SASP, but can also be linked with angiogenesis after infarction^34^. Furthermore, it is discussed as a biomarker for the risk and outcome for ischemic stroke^35,36^.

#### Cancer and Stroke

Co-morbidity has been described for stroke and pancreatic cancer^37^, based in part on cancer associated hypercoagulation. We added the four extra SASP factors from Valenzuela, et al. ^18^, their Table 1 (“Senescence-associated secretory phenotype (SASP) factors with potential effect on platelets aggregation and the fibrinolytic system”), that are not already included in the canonical map (that is, MMP2, FN1, THPO and CSF3). We then constructed a revised pathway map, which is mostly stable with respect to the canonical one except that clusters 1 and 2 are merged. In this revised pathway map, inconsistently regulated cell-cycle genes are thus forming the resulting top-left cluster, while the SASP factors are found upregulated in cluster 9, second row to the left. In the revised map, these include the upregulated IL6. Another upregulated SASP factor, IL1B, is found in cluster 19, third row. The other SASP factors involved in coagulation, including the ones that were added (red arrows), display only some moderate upregulation.

#### Mitochondrial dysfunction in mice

We recently contributed to an investigation of the effects of mitochondrial heteroplasmy in mice, based on a conplastic adenine-repeat variation (9 to 13A) in the origin of light-strand DNA replication of the mitochondrial genome, which causes shorter lifespan in female mice^38^. Overlaying the corresponding gene expression data onto the canonical senescence pathway map (Fig. 3, bottom) demonstrates no clear pattern except in cluster 2, where specifically the downstream genes (SIRT1, LMNB1, SERPINE1, TFDP2) are regulated as expected in cellular senescence; in fact, the SASP factor SERPINE1 (also known as PAI1) is the gene with the strongest upregulation in the entire pathway map.

**Fig. 3:**
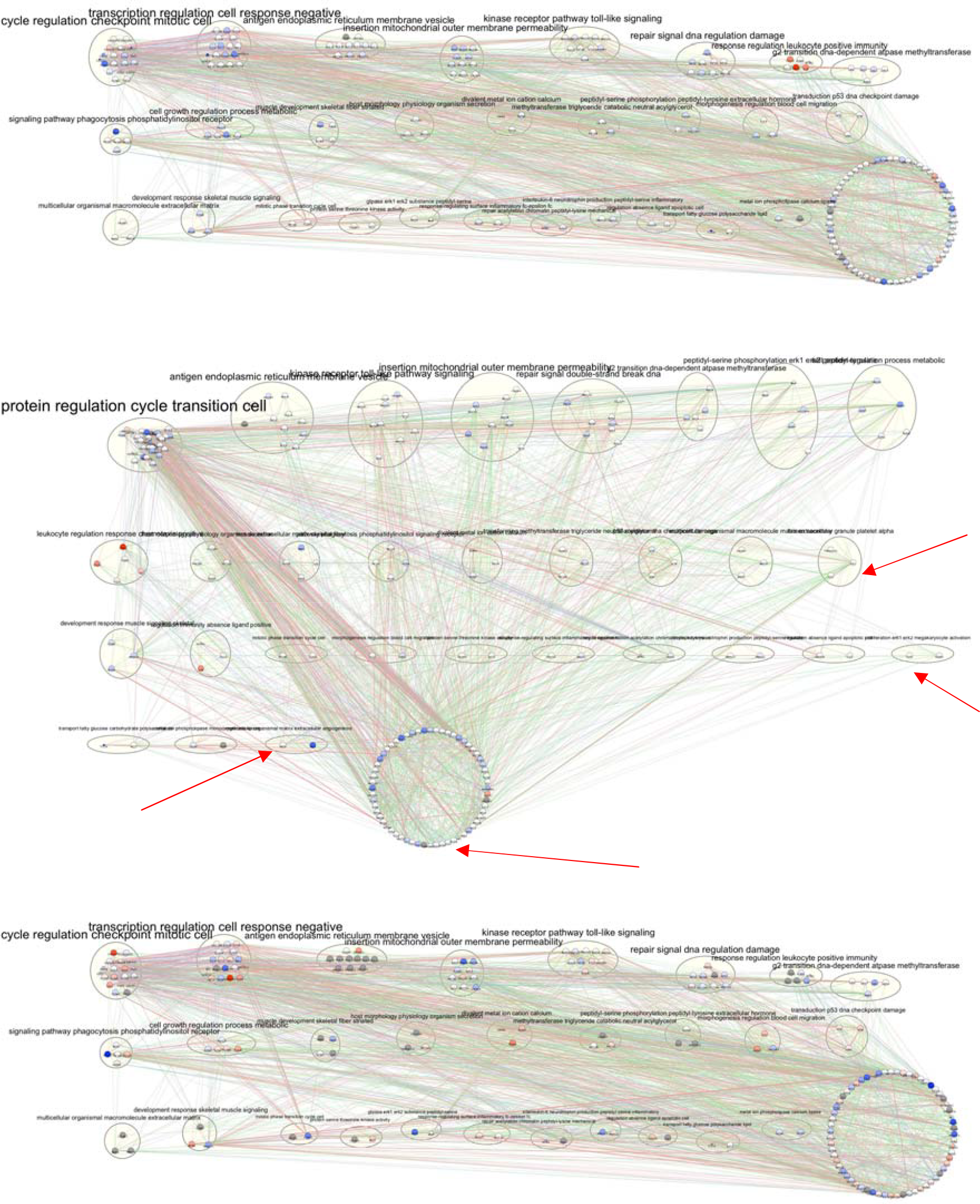
Senescence in stroke and mitochondrial dysfunction. The canonical senescence pathway map (top and bottom), based on the SASP and the KEGG cellular senescence genes, and an amended one (middle), adding four genes of interest and omitting any manual layout. Genes upregulated in disease or dysfunction (GSE22255, ref^33^, top and middle, Hirose et al.^38^ bottom) are shown in red, downregulated genes are shown in blue, and grey denotes genes for which no expression values are available. See Fig. 1 for further explanations.

## Discussion

The senescence pathway maps described here were generated in an automated fashion. Manual fine-tuning was only required for the initial selection of the underlying network data. This increases our confidence that the same approach can be followed by researchers who wish to obtain a quick overview of their high-throughput data in the context of their specific choice of network input genes. The approach also worked for our recent analyses of genes implicated in human and *C. elegans* health^39^. In all cases, public network data were used to generate a specific network based on a list of input genes by GeneMANIA^19^, which was then clustered into “pathways” by AutoAnnotate^20^, in turn employing ClusterMaker^40^. Here, the only fine-tuning came in, as follows. If a large amount of interactions is publicly available for the input genes, they form a tight “hairball” so that clustering by the default approaches offered by AutoAnnotate returns a single cluster, or no results at all. Thus, employing all the default network sources of GeneMANIA, clustering of the senescence genes that gave rise to the canonical *senescence pathway map* was then dominated by a single large cluster. Therefore, we limited the underlying network information to three of the six available sources (co-localization, genetic and protein interactions but not co-expression, pathways or shared domains) to obtain the map as presented in Fig. 1. In case of health genes based on genome-wide association studies, the default (larger) set of network sources could be used^39^ without resulting in a tight “hairball”, since many of the genes included there featured only sparse connectivity as is typical for a gene list based on genome-wide association.

The annotation of the clusters/pathways based on frequent words in the GO terms of the pathway genes was completely automated as well (by AutoAnnotate, in turn employing WordCloud). In the maps presented, we deviated slightly from the default WordCloud parameters, in setting the “max. number of words per cluster label” to the maximum of 5, to display as much information as possible, and by setting the “Adjacent word bonus” to 0; the default “Adjacent word bonus” of 8 triggered the listing of some rare words adjacent to frequent words, such as “vesicle-mediated transport” in case of the second-largest cluster, caused by a GO annotation of one gene (SERPIN1: “regulation of vesicle-mediated transport”). The layout (the determination of the exact positioning of the genes) was based on expert knowledge, but this step is optional, it pleases the eye (by avoiding label overlap) and enables easier understanding (by placing “upstream” and “downstream” genes into a presumably right order).

There are three major advantages of our approach, in comparison to the mapping of high-throughput data onto existing curated pathways such as the ones offered by KEGG or WikiPathways. For one, our approach works even if there is no expert pathway available that can be expected to help in understanding a high-throughput dataset. Also, our approach is not biased by the expert knowledge included in the structure of existing pathways. And there is flexibility in the selection of the gene list, here shown by assembling the canonical senescence gene list from two sources to start with, and later on by adding a few specific genes for application-specific investigations.

Our approach has limitations. In some scenarios it is possible that the pathway map defies common sense in grouping together genes whose interaction is based on the underlying network sources, but the genes are not known to work together based on the literature. It may be hard to decide whether such pathways are biologically meaningful, or whether these only reflect the noise in the underlying public interaction data. In our case, the canonical cell-cycle genes (CDKNs, CDKs; CCNs i.e. cyclins) were mostly, but not exclusively, distributed to the two largest clusters/pathways. Moreover, there are still a number of steps required for the recipe from the input gene list towards the pathway map, but these could be fully automated in principle. A final issue is the low dimensionality of any pathway map: in a 2D plane, and even in 3D space^1,41^, only the most prominent groups of genes can be visualized as genes that are “working together”. All the other groupings, many of which may act in parallel, or be realized only in some specific cellular contexts, are reflected in the large number of edges that connect the genes of the different clusters/pathways. Still, the plausibility of the resulting pathway maps does not necessarily come as a surprise. Already some time ago, Dutkowski, et al. ^42^ observed that the GO gene set hierarchy can, to some degree, be inferred from protein interaction data. Further, the experience of other researchers with GeneMANIA and AutoAnnotate includes many meaningful clusterings of networks as described in the original publications as well as in work citing these.

This work made intense use of the benefits that FAIR principles of scientific data already offer. Data were found and accessed with GEO and GEO2R, and they were interoperable via GeneMania, AutoAnnotate and Cytoscape, supporting reuse that enabled new insights into cellular phenotypes. This work shall be extended over time with data from more diseases to which senescence is considered to contribute. While the pathway maps themselves were generated in an automated fashion, the selection of experimental data was not. Thus, we have not yet reached the limits as to the degree of automation of data analyses, for senescence and other cellular phenotypes.

## Methods

### Construction of the canonical senescence pathway map

For the canonical senescence pathway map, we provided 33 known senescence markers assembled by Hernandez-Segura et al. ^4^ (Fig. S2 therein) together with the 160 genes included in the human “cellular senescence” KEGG pathway (see Suppl. File 1) to the Cytoscape application *GeneMANIA*^19^, version 3.4.1, downloaded October 2017. We used default settings except that we limited the underlying interaction data to co¬localization, genetic and protein interactions, to create a functional interaction network that is complemented with the GeneMANIA default of 20 connecting genes. (The overlap between the 33 markers from Hernandez-Segura et al and the 160 KEGG genes consisted of CDKN1A, CDKN2A, CXCL8 and IL6; including the 20 genes added by GeneMANIA, the senescence pathway map thus consists of 209 genes.) For clustering and annotation of the clusters based on the “annotation name” column of GO annotations collected by GeneMANIA, we used *AutoAnnotate*^20^ v1.2, downloaded October 2017, in Quick start modus to enable “layout network to prevent cluster overlap”, so that a map of disjoint clusters (senescence pathways) was generated, supplemented by a second advanced annotation step to increase the “max. number of words per cluster label” to the largest possible value of 5, and setting the “Adjacent word bonus” to 0. Cluster annotations were generated by AutoAnnotate based on these parameters using WordCloud^43^ v3.1.1, downloaded January 2018; using an adjacent word bonus results in cluster annotations with more words that do not feature an obvious relationship to cellular senescence. Thus, with two minor modifications, we conducted the same pathway mapping approach as Möller et al. ^39^. We performed a manual layout of the clusters/pathways guided by expert knowledge, placing genes considered “upstream” in senescence-related signaling on top, and placing genes considered “downstream” (specifically components of the SASP) further down.

### Overlaying of expression data onto pathway maps

We searched the GEO (Gene Expression Omnibus) database in July 2018 for datasets/series with the search term “cellular senescence”, limited to the type “Expression profiling by array”, sorting by the number of samples. From the resulting list, we selected GSE19899 as the second-largest dataset (50 samples); the largest dataset, GSE40489, was very specific to lymphoma and therefore ignored. The gene expression (GPL570) subset of the GSE19899 dataset consists of two experiments/batches (the latter one consisting of the last 6 samples, marked explicitly “E2”), and both experiments consist of two replicates, A and B. From experiment 1, we took the two “MLP (Growing)” replicates as *control*, and the two “MLP (Senescent)” replicates as *senescent* samples, ignoring quiescent samples as well as samples treated with shRNA (MLP samples are labeled as “Growing cells expressing a vector control”). From experiment 2, we took the two “(Growing)” replicates as *control*, and the two “(Senescent)” replicates as *senescent* samples. All samples are from human lung fibroblast IMR-90 cells, featuring control versus Ras-induced senescence.

We also investigated RNA-seq based datasets of cellular senescence. RNAseq data from one study^4^ was obtained from the Gene Expression Omnibus: Herranz et al. (accession GSE61130; we only contrasted senescent versus non-senescent cells; any manipulation related to ZFP36L1 was ignored). In addition, RNAseq data from Hernandez-Segura et al. was downloaded from Array Express (accession E-MTAB5403). Data was converted to log2-fold changes, from the normalized data.

To investigate senescence in a variety of disease scenarios, we selected the following disease datasets based on in-house expertise. Prostate cancer expression data^28^ (GSE46691) as prepared in a readily interpretable format^27^ was read into R version 3.5.1. Ensembl gene IDs were converted to HUGO gene IDs with biomaRt^44^, and mean expression of samples with Gleason score >= 7 minus mean expression levels of samples with a lower Gleason score was computed as logFC. For pancreatic cancer, we selected the only GEO hit with the search term ("cellular senescence" "pancreatic cancer"), GSE81368. We selected all 3 samples with growth state “non-senescent” as control, and all 6 samples with growth state “senescent”, independent of the experimental factors gender (male/female) and protocol (“Peroxide”/“Replication”). A wider search with the term (senescence "pancreatic cancer") retrieved one more dataset, that is, GSE28155. Here, we selected all three samples “BxPC3 cells transfected with control vector” as senescent, and all three samples “BxPC3 cells transfected with vector expressing shJMJD3” as control, since KDM6B (aka JMJD3) is a tumor-suppressive mediator of KRAS-induced senescence. For stroke, we found no GEO hit with the search term ("cellular senescence" "ischemic stroke”). Using the search term ("ischemic stroke" "peripheral blood mononuclear cells”), we took GSE22255, which has the largest number of samples (i.e. 40). We selected all 20 samples labeled “control” as control, and all 20 samples labeled “IS patient” for the senescent state, based on the “Characteristics” column. Finally, mapping liver gene expression data changes from the C57BL/6J-mtAKR/J strain of mice (compared to control C57BL/6J mice) that feature heteroplasmy-related dysfunction triggering a lower lifespan and a metabolic impairment^38^, we directly took the “effect size” column from their “Supplementary File S1”, and used the gene symbols for mapping directly from mouse to human.

For each GEO array-based dataset, we used the GEO2R tool^45^ to compute fold-changes using default parameters, inspected the “Values” boxplot, and downloaded the resulting table, imported it into *Numbers* (which works similar to Excel, but does not require changing “General” manually to “Text” column format for the gene names), and exported to CSV format. Selecting the “Gene.symbol” column as key column of the table and selecting the “gene name” column created by GeneMANIA as the “Key column for network”, we established matching gene names in the GEO2R and GeneMANIA tables as the common reference and then imported the tables into Cytoscape. In case of GSE81368 (pancreatic cancer), no gene symbols were available, but we could match the GB_ACC of GEO2R to the RefSeq mRNA ID in the GeneMANIA table, after removing all trailing version numbers („.1“, „.2“, „.3“, etc.) from the GB_ACC. Direct import of the text files from GEO2R is hindered by quotation marks, which are removed by *Numbers*. Finally, we created/copied and adjusted the “Style” of the resulting network so that the logFC values from GEO2R are mapped continuously to a red-blue color scale with the appropriate max/min settings, adding a handle to map a logFC of 0 to white.

The accompanying web presentation uses CytoscapeJS to present the pathways. Genes can be selected via their cluster or by the GeneOntology terms they are annotated with. Any such selection of genes is referenced to the MEM^46^ and g:Profiler^47^ web services.

## Acknowledgements

This project has received funding from the European Union’s Horizon 2020 research and innovation program under Grant agreement No 633589 (Aging with Elegans). This publication reflects only the authors’ views and the Commission is not responsible for any use that may be made of the information it contains.

## Competing interests

The authors declare that they have no competing interests.

## Authors’ contributions

Study design: GF. Collection of data: GF, SM, IB. Analysis of data: GF, SM, IB, AK. Manuscript writing and editing: GF, SM, AK, RJ, LH, HME, CJ, OH, MAW, BK, OW, SI, RK, FL, UW, MH. All authors reviewed and approved the final manuscript.

## Supplemental Information

Supplemental File 1. Excel Tables of all input genes and of all output clusters/pathways.

Supplemental File 2. Cytoscape (v. 3.4.1) file of all networks.

## References

1 Zhou, G. & Xia, J. OmicsNet: a web-based tool for creation and visual analysis of biological networks in 3D space. Nucleic acids research 46, W514-W522, doi:10.1093/nar/gky510 (2018).

2 Hayflick, L. & Moorhead, P. S. The serial cultivation of human diploid cell strains.Exp Cell Res 25, 585-621 (1961).

3 Hernandez-Segura, A., Nehme, J. & Demaria, M. Hallmarks of Cellular Senescence. Trends Cell Biol 28, 436-453, doi:10.1016/j.tcb.2018.02.001 (2018).

4 Hernandez-Segura, A. et al. Unmasking Transcriptional Heterogeneity in Senescent Cells. Current biology: CB 27, 2652-2660 e2654, doi:10.1016/j.cub.2017.07.033 (2017).

5 Yanai, H. & Fraifeld, V. E. The role of cellular senescence in aging through the prism of Koch-like criteria. Ageing research reviews 41, 18-33, doi:10.1016/j.arr.2017.10.004 (2018).

6 Demaria, M. et al. An essential role for senescent cells in optimal wound healing through secretion of PDGF-AA. Dev Cell 31, 722-733, doi:10.1016/j.devcel.2014.11.012 (2014).

7 Krizhanovsky, V. et al. Senescence of activated stellate cells limits liver fibrosis. Cell 134, 657-667, doi:10.1016/j.cell.2008.06.049 (2008).

8 Xu, M. et al. Senolytics improve physical function and increase lifespan in old age. Nat Med, doi:10.1038/s41591-018-0092-9 (2018).

9 Baker, D. J. et al. Naturally occurring p16-positive cells shorten healthy lifespan. Naturr 530, 184-189, doi:10.1038/nature16932 (2016).

10 Baar, M. P. et al. Targeted Apoptosis of Senescent Cells Restores Tissue Homeostasis in Response to Chemotoxicity and Aging. Cell 169, 132-147 e116, doi:10.1016/j.cell.2017.02.031 (2017).

11 NIH. (seer.cancer.gov, 2018).

12 NIH. (https://www.cancer.gov/types/pancreatic/hp/pancreatic-treatment-pdq, 2014).

13 Gonzalez-Meljem, J. M., Apps, J. R., Fraser, H. C. & Martinez-Barbera, J. P. Paracrine roles of cellular senescence in promoting tumourigenesis. Br J Cancer 118, 1283-1288, doi:10.1038/s41416-018-0066-1 (2018).

14 Moir, J. A., White, S. A. & Mann, J. Arrested development and the great escape—the role of cellular senescence in pancreatic cancer. Int J Biochem Cell Biol 57, 142-148, doi:10.1016/j.biocel.2014.10.018 (2014).

15 Childs, B. G. et al. Senescent cells: an emerging target for diseases of ageing. Nat Rev Drug Discov 16, 718-735, doi:10.1038/nrd.2017.116 (2017).

16 Hisada, Y. & Mackman, N. Cancer-associated pathways and biomarkers of venous thrombosis. Blood 130, 1499-1506, doi:10.1182/blood-2017-03-743211 (2017).

17 Posada-Duque, R. A., Barreto, G. E. & Cardona-Gomez, G. P. Protection after stroke: cellular effectors of neurovascular unit integrity. Front Cell Neurosci 8, 231, doi:10.3389/fncel.2014.00231 (2014).

18 Valenzuela, C. A., Quintanilla, R., Moore-Carrasco, R. & Brown, N. E. The Potential Role of Senescence As a Modulator of Platelets and Tumorigenesis. Frontiers in oncology 7, 188, doi:10.3389/fonc.2017.00188 (2017).

19 Zuberi, K. et al. GeneMANIA prediction server 2013 update. Nucleic acids research 41, W115-122, doi:10.1093/nar/gkt533 (2013).

20 Kucera, M., Isserlin, R., Arkhangorodsky, A. & Bader, G. D. AutoAnnotate: A Cytoscape app for summarizing networks with semantic annotations. F1000Research 5, 1717, doi:10.12688/f1000research.9090.1 (2016).

21 Kanehisa, M. & Goto, S. KEGG: kyoto encyclopedia of genes and genomes. Nucleic acids research 28, 27-30 (2000).

22 Chicas, A. et al. Dissecting the unique role of the retinoblastoma tumor suppressor during cellular senescence. Cancer cell 17, 376-387, doi:10.1016/j.ccr.2010.01.023 (2010).

23 Herranz, N. et al. mTOR regulates MAPKAPK2 translation to control the senescence-associated secretory phenotype. Nature cell biology 17, 1205-1217, doi:10.1038/ncb3225 (2015).

24 Sadasivam, S. & DeCaprio, J. A. The DREAM complex: master coordinator of cell cycle-dependent gene expression. Nat Rev Cancer 13, 585-595, doi:10.1038/nrc3556 (2013).

25 Chan, C. J., Andrews, D. M. & Smyth, M. J. Can NK cells be a therapeutic target in human cancer? European journal of immunology 38, 2964-2968, doi:10.1002/eji.200838764 (2008).

26 Lasry, A. & Ben-Neriah, Y. Senescence-associated inflammatory responses: aging and cancer perspectives. Trends in immunology 36, 217-228, doi:10.1016/j.it.2015.02.009 (2015).

27 Golightly, N. P., Bell, A., Bischoff, A. I., Hollingsworth, P. D. & Piccolo, S. R. Curated compendium of human transcriptional biomarker data. Scientific data 5, 180066, doi:10.1038/sdata.2018.66 (2018).

28 Erho, N. et al. Discovery and validation of a prostate cancer genomic classifier that predicts early metastasis following radical prostatectomy.PloS one 8, e66855, doi:10.1371/journal.pone.0066855 (2013).

29 R Core Team. (https://www.R-project.org/, 2018).

30 Kogan-Sakin, I. et al. Prostate stromal cells produce CXCL-1, CXCL-2, CXCL-3 and IL-8 in response to epithelia-secreted IL-1. Carcinogenesis 30, 698-705, doi:10.1093/carcin/bgp043 (2009).

31 Wang, T. et al. Senescent Carcinoma-Associated Fibroblasts Upregulate IL8 to Enhance Prometastatic Phenotypes. Molecular cancer research: MCR 15, 3-14, doi:10.1158/1541-7786.MCR-16-0192 (2017).

32 Yamamoto, K. et al. Loss of histone demethylase KDM6B enhances aggressiveness of pancreatic cancer through downregulation of C/EBPalpha. Carcinogenesis 35, 2404-2414, doi:10.1093/carcin/bgu136 (2014).

33 Krug, T. et al. TTC7B emerges as a novel risk factor for ischemic stroke through the convergence of several genome-wide approaches. Journal of cerebral blood flow and metabolism: official journal of the International Society of Cerebral Blood Flow and Metabolism 32, 1061-1072, doi:10.1038/jcbfm.2012.24 (2012).

34 Gertz, K. et al. Essential role of interleukin-6 in post-stroke angiogenesis. Brain 135, 1964-1980, doi:10.1093/brain/aws075 (2012).

35 Esenwa, C. C. & Elkind, M. S. Inflammatory risk factors, biomarkers and associated therapy in ischaemic stroke. Nat Rev Neurol 12, 594-604, doi:10.1038/nrneurol.2016.125 (2016).

36 Bustamante, A. et al. Prognostic value of blood interleukin-6 in the prediction of functional outcome after stroke: a systematic review and meta-analysis. J Neuroimmunol 274, 215-224, doi:10.1016/j.jneuroim.2014.07.015 (2014).

37 Bonnerot, M. et al. Cerebral ischemic events in patients with pancreatic cancer: A retrospective cohort study of 17 patients and a literature review. Medicine 95, e4009, doi:10.1097/MD.0000000000004009 (2016).

38 Hirose, M. et al. Low-level mitochondrial heteroplasmy modulates DNA replication, glucose metabolism and lifespan in mice. Sci Rep 8, 5872, doi:10.1038/s41598-018-24290-6 (2018).

39 Moeller, S. et al. Healthspan pathway maps in C. elegans and humans highlight transcription, prolifera-tion/biosynthesis and lipids. bioRxiv doi:10.1101/355131 (2018).

40 Morris, J. H. et al. clusterMaker: a multi-algorithm clustering plugin for Cytoscape. BMC bioinformatics 12, 436, doi:10.1186/1471-2105-12-436 (2011).

41 Gilbert, D. R., Schroeder, M. & van Helden, J. Interactive visualization and exploration of relationships between biological objects. Trends Biotechnol 18, 487-494 (2000).

42 Dutkowski, J. et al. A gene ontology inferred from molecular networks. Nature biotechnology 31, 38-45, doi:10.1038/nbt.2463 (2013).

43 Oesper, L., Merico, D., Isserlin, R. & Bader, G. D. WordCloud: a Cytoscape plugin to create a visual semantic summary of networks. Source code for biology and medicine 6, 7, doi:10.1186/1751-0473-6-7 (2011).

44 Durinck, S., Spellman, P. T., Birney, E. & Huber, W. Mapping identifiers for the integration of genomic datasets with the R/Bioconductor package biomaRt. Nature protocols 4, 1184-1191, doi:10.1038/nprot.2009.97 (2009).

45 Barrett, T. et al. NCBI GEO: archive for functional genomics data sets—update. Nucleic acids research 41, D991-995, doi:10.1093/nar/gks1193 (2013).

46 Adler, P. et al. Mining for coexpression across hundreds of datasets using novel rank aggregation and visualization methods. Genome Biol 10, R139, doi:10.1186/gb-2009-10-12-r139 (2009).

47 Reimand, J., Kull, M., Peterson, H., Hansen, J. & Vilo, J. g:Profiler—a web-based toolset for functional profiling of gene lists from large-scale experiments. Nucleic acids research 35, W193-200, doi:10.1093/nar/gkm226 (2007).

## Data Citations

Chicas A. et al., Gene Expression Omnibus GSE19899 (2010)

Herranz N. et al., Gene Expression Omnibus GSE61130 (2015)

Erho N. et al., Gene Expression Omnibus GSE46691 (2013)

Tsao M. et al., Gene Expression Omnibus GSE81368 (2017)

Yamamoto K. et al., Gene Expression Omnibus GSE28155 (2014)

Krug T. et al., Gene Expression Omnibus GSE22255 (2011)

Guryev V. ArrayExpress E-MTAB-5403 (2015)

Piccolo S. et al., Open Science Framework DOI:10.17605/OSF.IO/SSK3T (2016)

KEGG PATHWAY map04218 (2017)

